# Blue light induces neuronal-activity-regulated gene expression in the absence of optogenetic proteins

**DOI:** 10.1101/572370

**Authors:** Kelsey M. Tyssowski, Jesse M. Gray

## Abstract

Optogenetics is widely used to control diverse cellular functions with light, requiring experimenters to expose cells to bright light. Because extended exposure to visible light can be toxic to cells, it is important to characterize the effects of light stimulation on cellular function in the absence of exogenous optogenetic proteins. Here we exposed cultured mouse cortical neurons that did not express optogenetic proteins to several hours of flashing blue, red, or green light. We found that exposing neurons to as short as one hour of blue, but not red or green, light results in the induction of neuronal-activity-regulated genes without inducing neuronal activity. Our findings suggest blue light stimulation is ill-suited to long-term optogenetic experiments, especially those that measure transcription.

**Significance Statement:** Optogenetics is widely used to control cellular functions using light. In neuroscience, channelrhodopsins, exogenous light-sensitive channels, are used to achieve light-dependent control of neuronal firing. This optogenetic control of neuronal firing requires exposing neurons to high-powered light. We ask how this light exposure, in the absence of channelrhodopsin, affects the expression of neuronal-activity-regulated genes, i.e., the genes that are transcribed in response to neuronal stimuli. Surprisingly, we find that neurons without channelrhodopsin express neuronal-activity-regulated genes in response to as short as an hour of blue, but not red or green, light exposure. These findings suggest that experimenters wishing to achieve longer-term (an hour or more) optogenetic control over neuronal firing should avoid using systems that require blue light.

## Introduction

With the development of optogenetic technologies over the past decade (1,2), it has become increasingly common to expose biological samples to high-powered light. Optogenetics enables light-based control over diverse cellular functions–including neuronal firing (3), transcription (4,5), and cell signaling (1)–via exogenous proteins that are activated by specific wavelengths of light. Results of such experiments can be difficult to interpret if light by itself, in the absence of optogenetic proteins, affects cellular processes. Therefore, it is important to characterize how light exposure affects biological samples.

Sustained light exposure can form free radicals that affect cellular processes. In cell culture experiments, hours-long light exposure lowers cell viability via toxic oxidation and free radical formation in the media (6–10). Furthermore, *D. melanogaster* larvae and *C. elegans* are sensitive to free radicals that accumulate internally when the animals are exposed to visible light for hours to days (11–13). In both cell culture and animal studies, shorter (i.e., blue-er) wavelengths of light have greater negative effects than longer wavelengths. Despite the potential for side effects with blue light, the majority of optogenetic proteins, including channelrhodopsin (ChR2) and cryptochrome-2 (CRY-2) (1,2), are activated by blue light in the 450-500 nm range, although other optogenetic proteins are responsive to longer wavelengths of light (3,14,15). In neuroscience, blue light stimulation has been used with exogenous channelrhodopsins to to control neuronal firing for hours to days (16–19), time scales likely to result in light-induced toxicity. Indeed, exposing cultured neurons to 20 hours of blue light flashing at 1Hz reduces viability (20). However, less severe cellular changes that may occur separately from, or in early stages of, cell death have not been characterized.

Here we focused on assessing the role of light exposure on transcription. We were particularly interested in characterizing the effects of light on transcription in neurons because optogenetically-driven neuronal activity induces expression of activity-regulated genes, such as *Fos* (21). Therefore, optogenetics could be a useful tool to precisely control neuronal activation to study the resulting activity-regulated gene expression. Furthermore, optogenetics can be used to directly control transcription (4,5) and to control signaling pathways that regulate transcription (1), suggesting that experimenters may wish to measure transcriptional outputs of optogenetic experiments in many contexts.

We therefore asked whether neuronal transcription is affected by one to six hours of blue, red, or green light exposure. We chose light wavelengths that activate published channelrhodopsin variants (3,14,15) and timepoints relevant to activity-regulated gene induction (22,23). We found that cultured neurons without channelrhodopsin induced the activity-regulated genes *Fos, Npas4*, and *Bdnf* when exposed to one or six hours of blue light, but not when exposed to red or green light. Our findings suggest light by itself, in the absence of optogenetic proteins, can induce transcription. Therefore, experiments that measure transcription following long-term optogenetic stimulation should take precautions, such as using longer-wavelength light, to avoid experimental confounds from this light-induced transcription.

## Materials and Methods

### Cell culture

Corticies were dissected from embryonic day 16 (E16) CD1 or C57/Bl6 mouse embryos of mixed sex. They were dissociated with papain (Worthington, (L)(S)003126). 250,000 dissociated cells/well were plated on 48-well Lumos OptiClear plates (Axion), which have opaque well walls and had been coated overnight with poly-ornithine (30mg/mL, Sigma) and laminin (5ug/mL) in water and then washed once with PBS. Cultures were maintained at 37°C at 5% CO2 in BrainPhys media (StemCell Technologies) without phenol red supplemented with SM1 (StemCell Technologies) and penicillin/streptomycin (ThermoFisher).

### Light Stimulation

Light stimulation was done using the Lumos system programmed with AxIS software with power set at 100% (Axion Biosystems). According to the manufacturer, 100% power corresponds to 3.9mW/mm^2^ for blue light, 1.9mW/mm^2^ for green light, and 2.2mW/mm^2^ for red-orange light. These irradiance measurements were taken from the bottom of a well with no media (personal correspondance with Axion Biosystems). The wavelengths used were 475nm (blue), 530nm (green), and 612nm (red-orange). The temperature was maintained by putting the plate on a 37°C warming plate (Bel-Art). The CO2 was maintained at 5% throughout the duration of the recording using the base provided with the Axion Lumos system. Neurons were silenced with APV (100uM, Tocris) and NBQX (10uM, Tocris) at least 8h before stimulation to replicate conditions that would be used in optogenetic experiments. Light-exposed wells and wells left in the dark were on the same plate.

### Temperature Measurement

We measured temperature using a thermocouple (Omega, 5TC-TT-K-30-36) inserted into a well that was exposed to light stimulation. The temperature on the thermometer attached to the thermocouple was monitored at the start of the stimulation and at least once an hour for the duration of the light exposure (up to 6h).

### RNA extraction and qPCR

Immediatley following stimulation, samples were collected in Trizol (Invitrogen), and total RNA was extracted using the RNeasy mini kit (QIAGEN) with in-column DNase treatment (QIAGEN) according to the instructions of the manufacturer. The RNA was then converted to cDNA using the High Capacity cDNA Reverse Transcription kit (Applied Biosystems). For quantative PCR (qPCR), we used SsoFast Evagreen supermix (BioRad) with primers in Table 1. For qPCR analysis, we used a standard curve for each primer to convert Ct values into relative amounts of expression. We normalized our neuronal-activity-regulated gene values by values for the housekeeping gene *Gapdh* in order to control for any differences in the amount of cDNA in each reaction. *Gapdh* values were not significantly different between conditions (p*>*0.1, two-sided t-test). Furthermore, *Gapdh* mRNA is highly expressed and highly stable, making it less likely to be altered by small changes in transcription. Each biological replicate was from a different dissection on a different day. T-tests testing fold induction were performed on log fold change values from biological replicates testing the difference from a fold change of 1.

**Table 1.**
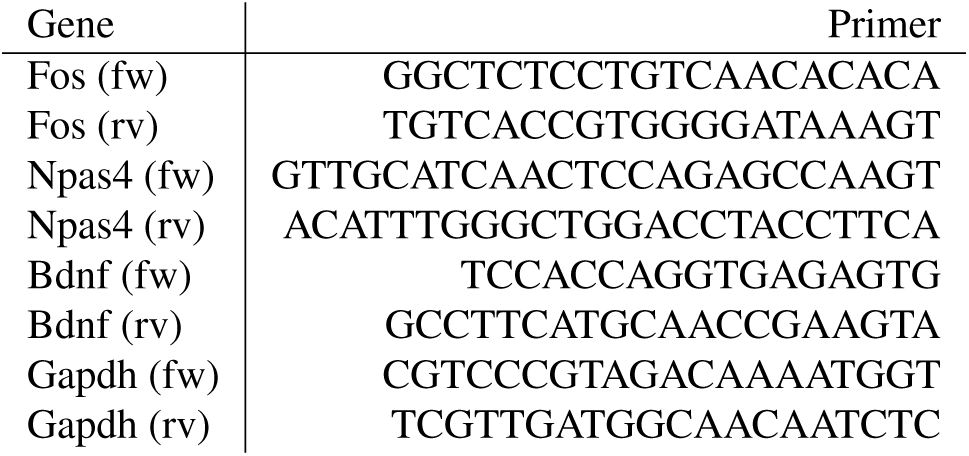
qPCR primers.

### Neuronal Activity Measurement

Neuronal activity was measured using neurons plated on Axion MEAs in 48-well optical plates. MEA plates were coated as above. Neurons from post-natal day 0 or 1 (P0-1) mice were dissociated and cultured as above. Recording was done using the Axion Maestro with the Axion software.

## Results

To determine the effect of light exposure on neurons, we exposed cultured cortical neurons that did not express channel-rhodopsin to a pattern of blue (475 nm) light consisting of 2-ms pulses at a frequency of 10 Hz. We used a light intensity of around 3.9 mW/mm^2^ (see Methods), which is similar to, or less than, the light intensity recommended for optogenetic activation of channelrhodopsin and similar molecules (3,14,15). After light exposure, we assessed the mRNA expression of the neuronal-activity-regulated gene *Fos* using qPCR. We found neurons exposed to just one hour of 10 Hz flashing blue light have 2.7-fold higher *Fos* mRNA expression than neurons left in the dark (Figure 1A). Following six hours of light exposure, we observed a four-fold induction of *Fos* mRNA, suggesting that blue light exposure–in the absence of optogenetic proteins–induces *Fos* mRNA expression.

**Fig. 1.**
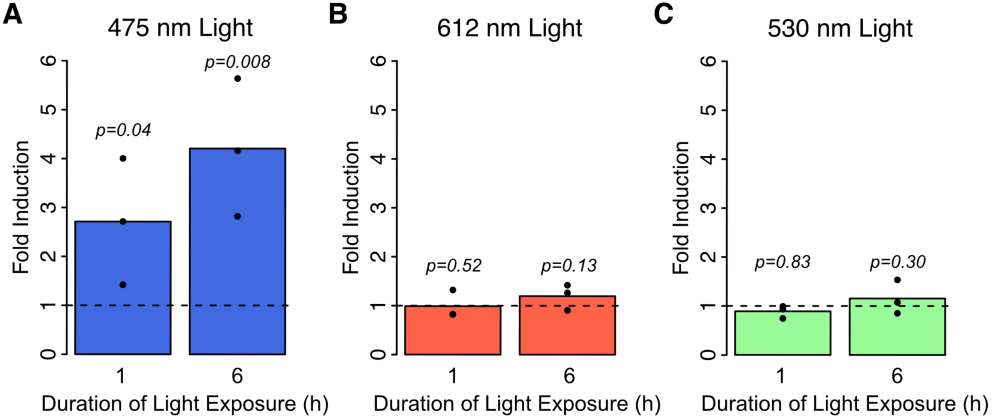
Cultured cortical neurons without channelrhodopsin were exposed to a pattern of 10Hz, 2-ms pulses of A) 475nm (blue), B) 612 nm (red), or C) 530nm (green) light for 1 or 6 hours. Induction of the activity regulated gene *Fos* was measured using RT-qPCR. Values plotted are the fold induction in mRNA expression at 6h compared to neurons not exposed to light. Bars represent the average of three biological replicates and dots are the values from each replicate. p-values from a one-sided Student’s t-test.

We next asked whether *Fos* is induced by red (612 nm) or green (530 nm) light. We exposed neurons to six hours of the same 10-Hz pattern and found that neither red nor green light exposure induces *Fos* expression (Figure 1B-C).

We then asked whether increasing the amount of blue light exposure increases gene induction. When we changed the pattern of blue-light stimulation to a frequency of 100 Hz, we found that neurons showed a 12-fold increase in *Fos* expression after six hours of light exposure. This was more than the four-fold increase we saw after six hours of exposure 10 Hz blue light (p=0.009, t-test), indicating that more light exposure results in more gene induction (Figure 2). However, we found that for red light, even the 100 Hz stimulation pattern failed to induce *Fos*, indicating that light-induced gene expression is specific to shorter (blue-er) wavelengths.

**Fig. 2.**
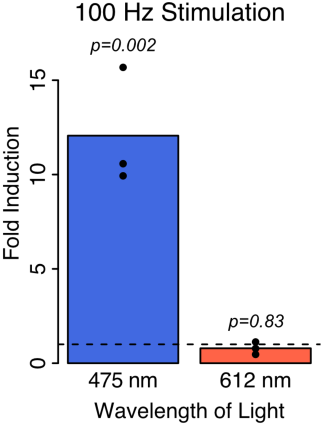
Cultured cortical neurons without channelrhodopsin were exposed to a pattern of 100Hz, 1-ms pulses of 475nm (blue) or 612 nm (red) light for 6 hours. Induction of the activity regulated gene *Fos* was measured using RT-qPCR. Values plotted are the fold induction in mRNA expression at 6h compared to neurons not exposed to light. Bars represent the average of three biological replicates and dots are the values from each replicate. p-values from a one-sided Student’s t-test.

We next investigated whether *Fos* induction might be due to one of several possible secondary effects of blue light exposure. *Fos* is usually induced in neurons as a result of the membrane depolarization that occurs during an action potential. However, neurons without channelrhodopsin grown on multi-electrode arrays did not fire more action potentials when exposed to light (Figure 3A), ruling out membrane depolarization as a cause of *Fos* induction. We also confirmed that the sustained blue light exposure did not substantially alter the temperature of the media. Even with 100Hz stimulation, the media remained at 37+/-1 °C for the duration of the experiment (Figure 3B). These findings suggest that the blue-light-induced gene expression is likely a direct rather than secondary effect of light exposure.

**Fig. 3.**
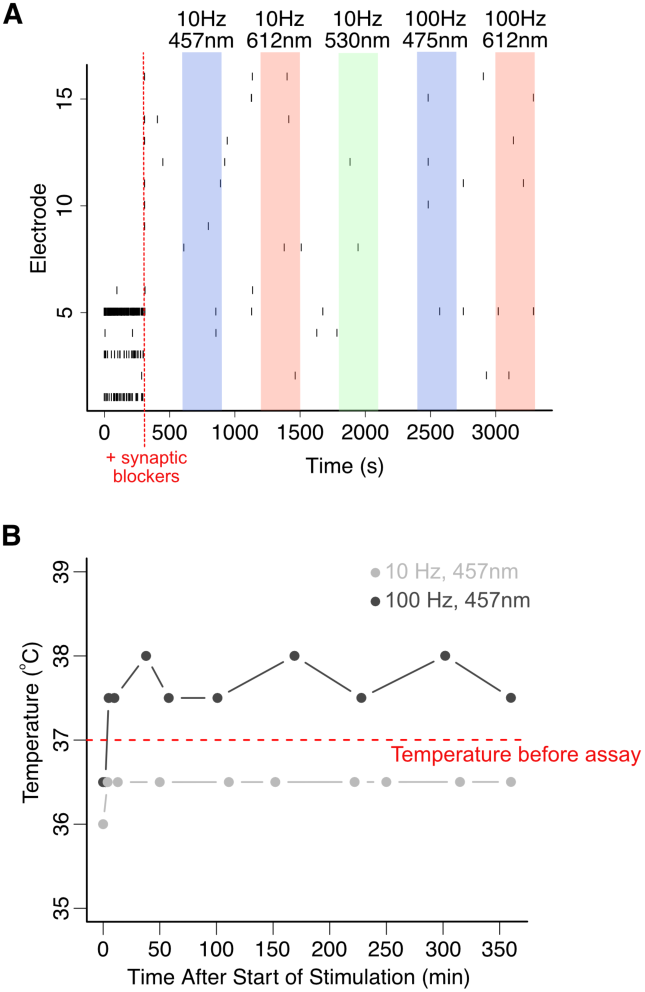
A) Cultured cortical neurons without channelrhodopsin plated on multi-electrode arrays were exposed to the indicated light conditions. As in all experiments, neurons were silenced before light exposure with synaptic blockers APV and NBQX. Each line represents an action potential. Light is ON at the highlighted times. Representative example from one experiment. B) Temperature measurements were taken from a well exposed to blue light at several time points during the course of a 6-hour experiment. The red line represents the temperature at which the neurons were cultured before starting the assay. Representative example from one experiment.

Finally, we asked whether others of the hundreds of neuronal-activity-regulated genes are induced by light exposure. Specifically, we assessed expression of *Bdnf* and *Npas4* mRNA using qPCR. We hypothesized that since *Bdnf* is regulated differently from *Fos* (22,23), it may not be induced by the *Fos*-regulating signaling pathways activated by blue light exposure. However, we found that *Bdnf* mRNA expression is induced by a six-hour exposure to blue, but not red or green, light (Figure 4A). Induction of *Npas4* mRNA, unlike *Fos* (24), is relatively specific to activated neurons (25). We thus reasoned that if the gene induction in response to blue light stimulation were activated as part of a response to oxidation and cell death (6–8), a neuronal-activation-specific gene might not be induced. Surprisingly, we found that a six-hour exposure to blue, but not red or green, light also resulted in induction of *Npas4* mRNA (Figure 4B). We therefore suspect that many neuronal-activity-regulated genes are induced by blue light exposure.

**Fig. 4.**
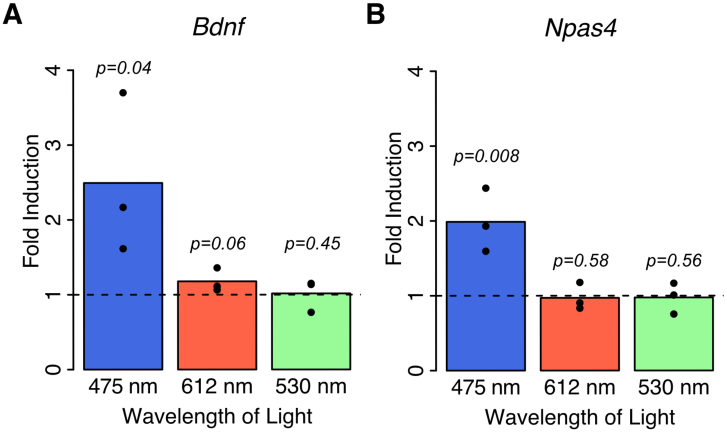
Cultured cortical neurons without channelrhodopsin were exposed to a pattern of 10Hz, 2-ms pulses of 475nm (blue), 612 nm (red), or 530nm (green) light for 6 hours. Induction of the activity regulated genes A) *Bdnf* and B) *Npas4* were measured using RT-qPCR. Values plotted are the fold induction in mRNA expression at 6h compared to neurons not exposed to light. Bars represent the average of three biological replicates and dots are the values from each replicate. p-values from a one-sided Student’s t-test.

## Discussion

We show in cultured cortical neurons without channel-rhodopsin, that extended exposure to blue light induces neuronal-activity-regulated genes. This gene induction does not occur in response to exposure to red or green light. Our findings indicate that blue light is ill-suited to optogenetic experiments that use long-term light exposure and those that assess changes in transcription in response to optogenetic stimulation. This work also emphasizes the importance of including experimental controls (26) in optogenetic experiments to determine the effects of light on cells in the absence of light-activated proteins.

Our finding that blue, but not red or green, light induces spurious transcription is consistent with other work demonstrating detrimental effects of short-wavelength light (8,20,27,28). Several studies that have compared the effects of blue light to other wavelengths of light both *in vitro* and in *C. elegans* have found that blue light has greater effects on cell viability (10), *C. elegans* behavior (11), and *C. elegans* survival (12). These data suggest that using optogenetic proteins that are activated by longer wavelengths of light (14,15) might allow experimenters to avoid side effects of light exposure.

We speculate that neurons induce transcription in response to blue light due to the oxidation that occurs in biological liquids in response to extended light exposure (6–9,20). Oxidative stress can induce transcription of primary response genes, including *Fos*, in a variety of cell types via activation of cell-signaling pathways, including the MAPK and NFkB path-ways (29). Because oxidative stress activates similar pathways as neuronal activity (22), we might expect oxidative stress to activate many neuronal-activity regulated genes. Indeed, we observed that blue light exposure induces all three of the neuronal-activity-regulated genes that we tested.

In neuronal cell culture systems, blue light exposure likely induces oxidation due to the presence of compounds such as riboflavin, tryptophan, and HEPES in cell culture media (28, 30–32). BrainPhys, the media used in this study, contains both riboflavin and HEPES (33), as does the common neuronal culture media, Neurobasal (manufacturer’s pamphlet). Therefore, supplementing neuronal culture media with antioxidants (9,16) or altering it to exclude compounds that cause oxidation (20) may mitigate the detrimental effects of blue light in culture systems. Alternatively, sensitive channelrhodopsins (21) can be used to minimize the duration of light exposure and thus its negative effects. Notably, we suspect that blue light exposure induces transcription in non-neural cultures, as the common cell culture media DMEM also contains riboflavin and HEPES (manufacturer’s pamphlet). Thus, spurious blue-light-induced gene expression may be a concern in any experiment that measures transcription in response to an opotogenetic stimulus, including those that use optogenetics to directly induce transcription in non-neural cells (4,5).

The toxic oxidation that occurs in culture media suggests that *in vitro* experiments may be particularly sensitive to blue light exposure. However, oxidation-prone compounds exist within cells and in interstitial fluids, suggesting that light exposure could also affect cells *in vivo*. Consistent with this idea, exposing *C. elegans* to blue light likely produces free radicals within the worm (11), and *C. elegans*, planeria, and *D. melanogaster* have free-radical-detecting cells that respond to light exposure in the absence of cell culture media (11,13,34). Alternatively, it is possible that endogenous opsins or cytochromes, which are expressed in our cultures (23) and in the brain (35), play a role in the observed gene induction, in which case we would expect to observe similar activity-regulated gene induction *in vivo*. Indeed, there is some evidence that blue light stimulation in the absence of channelrhodopsin may induce *Fos* expression in the rat brain (36), although it is not clear whether this is due to light stimulation or other factors, such as the trauma from implanting the optical fiber. Therefore, it will be important for future work to assess the impact of blue light exposure on neuronal transcription *in vivo*.

## ACKNOWLEDGEMENTS

KMT received funding from the NSF Graduate Research Fellowship Program DGE1144152, DEG1745303. JMG’s lab is supported by NIHMH116223, the Harvard-Armenise Foundation, and the Kaneb family. We thank Ricardo Henriques for the LaTeX bioRxiv paper template.

## AUTHOR CONTRIBUTIONS

KMT conceptualized project, designed research, performed research, analyzed data, and wrote the paper. JMG designed research and edited the paper.

## Bibliography

1. Hannes M. Beyer, Sebastian Naumann, Wilfried Weber, and Gerald Radziwill. Optogenetic control of signaling in mammalian cells. Biotechnology Journal, 10(2):273–283, 2015. ISSN 18607314. doi: 10.1002/biot.201400077.

2. Edward S Boyden, Feng Zhang, Ernst Bamberg, Georg Nagel, and Karl Deisseroth. Millisecond-timescale, genetically targeted optical control of neural activity. Nature neuroscience, 8(9):1263–8, 9 2005. ISSN 1097-6256. doi: 10.1038/nn1525.

3. John Y Lin. A user’s guide to channelrhodopsin variants: features, limitations and future developments. Experimental physiology, 96(1):19–25, 1 2011. ISSN 1469-445X. doi: 10. 1113/expphysiol.2009.051961.

4. Lauren R. Polstein and Charles A. Gersbach. A light-inducible CRISPR-Cas9 system for control of endogenous gene activation. Nature Chemical Biology, 11(3):198–200, 2015. ISSN 15524469. doi: 10.1038/nchembio.1753.

5. Yuta Nihongaki, Shun Yamamoto, Fuun Kawano, Hideyuki Suzuki, and Moritoshi Sato. CRISPR-Cas9-based photoactivatable transcription system. Chemistry and Biology, 22(2): 169–174, 2015. ISSN 10745521. doi: 10.1016/j.chembiol.2014.12.011.

6. Harold F Blum. Photodynamic Action. Physiological Reviews, 12(1):23–55, 1932.

7. Arthur Richardson. The action of light in preventing putrefactive decomposition; and in inducing the formation of hydrogen peroxide in organic liquids. J. Chem. Soc. Trans., 63: 1109–1130, 1893.

8. J D Stoien and R J Wang. Effect of near-ultraviolet and visible light on mammalian cells in culture II. Formation of toxic photoproducts in tissue culture medium by blacklight. Proceedings of the National Academy of Sciences of the United States of America, 71(10):3961–5, 1974. ISSN 0027-8424. doi: 10.1073/pnas.71.10.3961.

9. Ram Dixit and Richard Cyr. Cell damage and reactive oxygen species production induced by fluorescence microscopy: effect on mitosis and guidelines for non-invasive fluorescence microscopy. The Plant Journal, 36(2):280–290, 2003. ISSN 09607412. doi: 10.1046/j. 1365-313X.2003.01868.x.

10. Sina Waldchen, Julian Lehmann, Teresa Klein, Sebastian Van De Linde, and Markus Sauer. Light-induced cell damage in live-cell super-resolution microscopy. Scientific Reports, 5:1–12, 2015. ISSN 20452322. doi: 10.1038/srep15348.

11. Nikhil Bhatla and H. Robert Horvitz. Light and Hydrogen Peroxide Inhibit C.elegans Feeding through Gustatory Receptor Orthologs and Pharyngeal Neurons. Neuron, 85(4):804–818, 2015. ISSN 10974199. doi: 10.1016/j.neuron.2014.12.061.

12. C. Daniel De Magalhaes Filho, Brian Henriquez, Nicole E. Seah, Ronald M. Evans, Louis R. Lapierre, and Andrew Dillin. Visible light reduces C. elegans longevity. Nature Communications, 9(1), 2018. ISSN 20411723. doi: 10.1038/s41467-018-02934-5.

13. Ananya R. Guntur, Pengyu Gu, Kendra Takle, Jingyi Chen, Yang Xiang, and Chung-Hui Yang. Drosophila TRPA1 isoforms detect UV light via photochemical production of H2O2. Proceedings of the National Academy of Sciences, 112(42):E5753–E5761, 2015. ISSN 0027-8424. doi: 10.1073/pnas.1514862112.

14. John Y Lin, Per Magne Knutsen, Arnaud Muller, David Kleinfeld, and Roger Y Tsien. ReaChR: a red-shifted variant of channelrhodopsin enables deep transcranial optogenetic excitation. Nature neuroscience, 16(10):1499–508, 10 2013. ISSN 1546-1726. doi: 10.1038/nn.3502.

15. Nathan C Klapoetke, Yasunobu Murata, Sung Soo Kim, Stefan R Pulver, Amanda Birdsey-Benson, Yong Ku Cho, Tania K Morimoto, Amy S Chuong, Eric J Carpenter, Zhijian Tian, Jun Wang, Yinlong Xie, Zhixiang Yan, Yong Zhang, Brian Y Chow, Barbara Surek, Michael Melkonian, Vivek Jayaraman, Martha Constantine-Paton, Gane Ka-Shu Wong, and Edward S Boyden. Independent optical excitation of distinct neural populations. Nature methods, 11(3):338–46, 3 2014. ISSN 1548-7105. doi: 10.1038/nmeth.2836.

16. Matthew S Grubb and Juan Burrone. Activity-dependent relocation of the axon initial segment fine-tunes neuronal excitability. Nature, 465(7301):1070–4, 6 2010. ISSN 1476-4687. doi: 10.1038/nature09160.

17. Ming Fai Fong, Jonathan P. Newman, Steve M. Potter, and Peter Wenner. Upward synaptic scaling is dependent on neurotransmission rather than spiking. Nature Communications, 6: 1–11, 2015. ISSN 20411723. doi: 10.1038/ncomms7339.

18. Carleton P Goold and Roger a Nicoll. Single-cell optogenetic excitation drives homeostatic synaptic depression. Neuron, 68(3):512–28, 11 2010. ISSN 1097-4199. doi: 10.1016/j.neuron.2010.09.020.

19. Seongjun Park, Ryan A. Koppes, Ulrich P. Froriep, Xiaoting Jia, Anil Kumar H. Achyuta, Bryan L. McLaughlin, and Polina Anikeeva. Optogenetic control of nerve growth. Scientific Reports, 5:1–9, 2015. ISSN 20452322. doi: 10.1038/srep09669.

20. John H. Stockley, Kimberley Evans, Moritz Matthey, Katrin Volbracht, Sylvia Agathou, Jana Mukanowa, Juan Burrone, and Ragnhildur T. Káradóttir. Surpassing light-induced cell damage in vitro with novel cell culture media. Scientific Reports, 7(1):1–11, 2017. ISSN 20452322. doi: 10.1038/s41598-017-00829-x.

21. Philipp Schoenenberger, Daniela Gerosa, and Thomas G Oertner. Temporal control of immediate early gene induction by light. PloS one, 4(12):e8185, 1 2009. ISSN 1932-6203. doi: 10.1371/journal.pone.0008185.

22. Anne E West and Michael E Greenberg. Neuronal Activity – Regulated Gene Transcription in Synapse Development and Cognitive Function. pages 1–21, 2011.

23. Kelsey M. Tyssowski, Nicholas R. DeStefino, Jin Hyung Cho, Carissa J. Dunn, Robert G. Poston, Crista E. Carty, Richard D. Jones, Sarah M. Chang, Palmyra Romeo, Mary K. Wurzelmann, James M. Ward, Mark L. Andermann, Ramendra N. Saha, Serena M. Dudek, and Jesse M. Gray. Different Neuronal Activity Patterns Induce Different Gene Expression Programs. Neuron, 98(3):530–546, 2018. ISSN 10974199. doi: 10.1016/j.neuron.2018.04. 001.

24. Trent Fowler, Ranjan Sen, and Ananda L. Roy. Regulation of primary response genes. Molecular Cell, 44(3):348–360, 2011. ISSN 10972765. doi: 10.1016/j.molcel.2011.09.014.

25. Yingxi Lin, Brenda L Bloodgood, Jessica L Hauser, Ariya D Lapan, Alex C Koon, Tae-Kyung Kim, Linda S Hu, Athar N Malik, and Michael E Greenberg. Activity-dependent regulation of inhibitory synapse development by Npas4. Nature, 455(7217):1198–204, 2008. ISSN 1476-4687. doi: 10.1038/nature07319.

26. Brian D. Allen, Annabelle C. Singer, and Edward S. Boyden. Principles of designing interpretable optogenetic behavior experiments. Learning and Memory, 22(4):232–238, 2015. ISSN 15495485. doi: 10.1101/lm.038026.114.

27. Kevin P. Cheng, Elizabeth A. Kiernan, Kevin W. Eliceiri, Justin C. Williams, and Jyoti J. Watters. Blue Light Modulates Murine Microglial Gene Expression in the Absence of Optogenetic Protein Expression. Scientific Reports, 6(January):1–11, 2016. ISSN 20452322. doi: 10.1038/srep21172.

28. Bernard F. Godley, Farrukh A. Shamsi, Fong-Qi Liang, Stuart G. Jarrett, Sallyanne Davies, and Mike Boulton. Blue Light Induces Mitochondrial DNA Damage and Free Radical Production in Epithelial Cells. Journal of Biological Chemistry, 280(22):21061–21066, 2005. ISSN 0021-9258. doi: 10.1074/jbc.M502194200.

29. RG Allen and M Tresini. OXIDATIVE STRESS AND GENE REGULATION. Free Radical Biology and Medicine, 28(3):463–499, 2000.

30. A. M. Edwards, E. Silva, B. Jofré, M. I. Becker, and A. E. De Ioannes. Visible light effects on tumoral cells in a culture medium enriched with tryptophan and riboflavin. Journal of Photochemistry and Photobiology, B: Biology, 24(3):179–186, 1994. ISSN 10111344. doi: 10.1016/1011-1344(94)07020-2.

31. Jose Luis Lepe-Zuniga, J. S. Zigler, and Igal Gery. Toxicity of light-exposed Hepes media. Journal of Immunological Methods, 103(1):145, 1987. ISSN 00221759. doi: 10.1016/0022-1759(87)90253-5.

32. G T Spierenburg, F T J J Oerlemans, J P R M van Laarhoven, and C H M M de Bruyn. Phototoxicity of N-2-Hydroxyethylpiperazine-N-2-ethanesulfonic Acid-buffered Culture Media for Human Leukemic Cell lines. Cancer Research, 44:2253–2254, 1984.

33. Fred H Gage and Cedric Bardy. Media compositions for neuronal cell culture, 2014.

34. Taylor R. Birkholz and Wendy S. Beane. The planarian *TRPA1* homolog mediates extraocular behavioral responses to near-ultraviolet light. The Journal of Experimental Biology, 220(14):2616–2625, 2017. ISSN 0022-0949. doi: 10.1242/jeb.152298.

35. Stuart N. Peirson, Stephanie Haiford, and Russell G. Foster. The evolution of irradiance detection: Melanopsin and the non-visual opsins. Philosophical Transactions of the Royal Society B: Biological Sciences, 364(1531):2849–2865, 2009. ISSN 14712970. doi: 10.1098/rstb.2009.0050.

36. Franz R. Villaruel, Franca Lacroix, Christian Sanio, Daniel W. Sparks, C. Andrew Chapman, and Nadia Chaudhri. Optogenetic activation of the infralimbic cortex suppresses the return of appetitive pavlovian-conditioned responding following extinction. Cerebral Cortex, 28(12): 4210–4221, 2018. ISSN 14602199. doi: 10.1093/cercor/bhx275.

